# STN1 upregulation promotes PARPi resistance in BRCA2-deficient cancer cells via replication fork protection and suppression of ssDNA gap formation

**DOI:** 10.64898/2026.05.15.725290

**Authors:** Zubair Ahmed Laghari, Na Li, Serdar Gayybov, Qi-En Wang, Weihang Chai

## Abstract

PARPi are effective therapy for BRCA1/2 mutant cancers, yet recurrent PARPi resistance frequently develops. The underlying mechanism of PARPi resistance remains largely unresolved. Here, we identify STN1, a component of the CTC1/STN1/TEN1 (CST) complex, as a modulator of PARPi resistance in BRCA2-deficient cells. RNA-seq analysis of PARPi-resistant cancer cells from BRCA2-mutated backgrounds shows largely distinct transcriptomic profiles with limited overlap, suggesting multiple routes to resistance. Notably, STN1 is consistently upregulated in resistant cells. We observe that overexpression of STN1 enhances Olaparib resistance in multiple BRCA2-deficient cell lines and alleviates DNA damage under replication stress. Mechanistically, we find that STN1 overexpression increases RAD51 loading to stalled replication forks while restricting MRE11 recruitment in BRCA2-deficient cells, thereby protecting stalled forks from nascent-strand degradation. Furthermore, STN1 overexpression rescues the accumulation of ssDNA gaps, a major determinant of PARPi sensitivity in BRCA2-deficient cells. Taken together, these findings suggest that elevated STN1 levels can partially compensate for BRCA2 loss by stabilizing stalled replication forks and limiting ssDNA gap accumulation. Our study uncovers a STN1-dependent pathway of replication stress tolerance that promotes PARPi resistance independently of homologous recombination restoration, highlighting STN1 as a potential biomarker and mechanistic contributor to therapeutic resistance in BRCA2-mutated cancers.

## Introduction

Poly(ADP-ribose) polymerase inhibitors (PARPi) have emerged as an effective targeted therapy for tumors with defects in homologous recombination repair (HRR) (1–5). The most frequently altered HRR genes are *BRCA1* and *BRCA2*, although pathogenic alterations in other HRR-associated genes, including PALB2, BARD1, BRIP1, RAD51C, and RAD51D, can also confer HRR deficiency in subsets of tumors (4,6). Clinically approved PARPi include Olaparib, Niraparib, Rucaparib, Talazoparib, and most recently Saruparib. These agents primarily target PARP1, with varying activity against PARP2, thereby disrupting PARP-dependent DNA damage responses. Inhibition of PARP catalytic activity prevents PARylation and the efficient resolution of DNA lesions, while PARP trapping stabilizes PARP-DNA complexes that impede replication fork progression and promote replication-associated DNA damage. Increasing evidence indicates that persistent single-stranded DNA (ssDNA) replication gaps are a major determinant of PARPi sensitivity and contribute to synthetic lethality independently of, and in addition to, replication-associated double-strand breaks (DSBs) (6–9). In HR-deficient cells, the inability to repair these replication-associated lesions results in genomic instability and ultimately cell death (6). Exploiting this synthetic lethal interaction between PARP inhibition and HRR deficiency, PARPi have been approved for the treatment of ovarian, breast, prostate, and pancreatic cancers harboring BRCA1/2 mutations or other biomarkers of HRR deficiency (6,10,11).

Despite significant clinical benefit of PARPi, therapeutic resistance frequently develops, leading to disease relapses or progression and posting a major challenge to patient survival (12,13). Accumulating evidence indicates that PARPi resistance arises through diverse genetic and non-genetic mechanisms that may include reversion mutations restoring BRCA function, epigenetic reactivation of silenced genes, restoration of HR, loss of PARP1 expression or mutations that diminish PARP trapping, and increased drug efflux (13–17). In addition, tumor cells can evade PARPi-induced cytotoxicity by protecting stalled replication forks, thereby limiting replication-associated DNA damage, or by reducing the formation or persistence of toxic PARP-DNA complexes (10,15).

BRCA1 and BRCA2 play distinct and essential roles in maintaining genome integrity (18,19). BRCA1 primarily promotes DNA end resection at DSBs during the initiation of HR (20), whereas BRCA2 facilitates RAD51 loading onto ssDNA to complete HR repair (21,22). Beyond HR, BRCA2 protects stalled replication forks by promoting RAD51 loading, suppresses ssDNA gap formation, and preventing MRE11-mediated degradation of nascent DNA (6,8,21–26). Restoration of BRCA1/2 through either secondary mutations or other mechanisms that re-establish HRR represents a major cause of acquired PARPi resistance (15,17,27–29). Additional resistance mechanisms involve alterations in DNA damage response (DDR) signaling, replication fork protection, and checkpoint pathways (30,31). To counteract such resistance, recent therapeutic strategies aim at disrupting these compensatory DDR pathways to restore PARPi sensitivity (30,32–34).

The CST complex, comprised of CTC1, STN1 and TEN1, is an oligonucleotide/oligosaccharide-binding fold (OB-fold)-containing ssDNA-binding that binds to ssDNA and ss/dsDNA junctions (35,36). Initially characterized for its role in telomere maintenance, CST has emerged as an important regulator of genome stability by promoting fork protection and modulating DSB repair (37–39). At telomeres, CST facilitates DNA polymerase α (POLα)-mediated C-strand fill-in and limits telomerase activity through the POT1-TPP1 complex to prevent G-strands overextension (40–42). Upon fork stalling, CST localizes at stalled forks, where it facilities RAD51 loading and protects nascent DNA from MRE11-dependent degradation (43,44). Accordingly, CST deficiency leads to ssDNA accumulation, increased chromosomal aberrations, and accumulation of DNA damage (44–46). At DSBs, CST functions downstream of 53BP1-Shieldin to restrict end resection, thereby promoting non-homologous end joining (NHEJ) while inhibiting HR (39,47).

The role of CST in PARPi response has been studied in BRCA1-deficient cells. Because CST limits DSB end resection through inhibition of EXO1 and BLM-DNA2 and promotes POLα-dependent fill-in of resected ends, CST loss restores end resection and HR-like repair in BRCA1-deficient cells, resulting in PARPi resistance (47,48). In contrast, CST depletion in BRCA2-deficient cells exacerbates genome instability and impairs cell viability in the absence of PARPi, indicating that CST and BRCA2 cooperate in replication fork protection rather than acting antagonistically (43).

Here, we investigated the mechanisms underlying PARPi resistance in BRCA2-deficient cancer cells. We compared the transcriptomes of the parental PARPi-sensitive BRCA2-deficient PEO1 cell line with those of two PARPi-resistant derivatives: PEO4, which arose from the same patient, and PEO1-R, which we established from PEO1 in this study. We found that PEO1-R and PEO4 exhibited largely distinct transcriptomic profiles, indicating that PARPi resistance can arise through multiple molecular routes rather than a single, uniform transcriptional program. Interestingly, STN1 was consistently upregulated in both resistant cell lines. We found that ectopic overexpression of STN1 in BRCA2-deficient cancer cells conferred increased resistance to Olaparib. Mechanistically, STN1 overexpression attenuated BRCA2 deficiency-induced DNA damage, restored RAD51 loading at stalled replication forks, and limited MRE11 access to stalled forks, and suppressed nascent strand degradation. In addition, STN1 overexpression diminished ssDNA gap formation in BRCA2-deficient cells. Together, our findings identify STN1 as a key mediator of replication fork protection that compensates for BRCA2 loss and promotes PARPi resistance, thereby uncovering a previously unrecognized mechanism of acquired PARPi resistance in BRCA2-deficient cancers.

## Materials and methods

### Cell culture

HeLa and MDA-MB-231 cells were obtained from the American Type Culture Collection (ATCC) and cultured in Dulbecco’s modified Eagle’s medium (DMEM, Cytiva HyClone) supplemented with 10% Cosmic Calf^TM^ Serum (cat# SH3008704, Cytiva HyClone). PEO1 was obtained from Millipore Sigma and was cultured in RPMI 1640 medium (cat# R8758, Millipore Sigma) supplemented with 10% fetal bovine serum (FBS, Atlanta Biologicals). PEO4 was a gift from Dr. Kenneth P. Nephew. HeLa BRCA2 knockout (KO) cells have been described previously (49). PEO1 cells stably expressing FLAG-STN1 and Myc-CTC1 were generated by retroviral transduction followed by selection with puromycin using previously described protocol (44,50). All cell lines used in this study were maintained at 37 °C in a humidified incubator supplied with 5% CO2.

### Development of PEO1-R

Olaparib-resistant PEO1-R cells were established from the parental PEO1 line through intermittent, incremental in vitro exposure to Olaparib. The drug concentration was gradually increased from 0.5 to 20 µM over a six-month period. At each concentration, cells were treated with Olaparib for 48 hours, after which the medium was replaced with drug-free complete medium. The cells were then allowed to recover and proliferate until reaching approximately 80-90% confluence. Once stable growth was maintained for at least one to two passages at each concentration, the Olaparib dose was doubled, and the treatment cycle was repeated until the final concentration was achieved.

### RNA sequencing and data analysis

Bulk RNA sequencing was performed on three independent biological replicates for each cell line. Total RNA was extracted using the RNeasy Mini Kit (Qiagen) following the manufacturer’s protocol. RNA integrity was assessed using an Agilent 2100 Bioanalyzer, and samples with RNA integrity number (RIN) ≥ 9 were used for library preparation. Poly(A)+ RNA enrichment, sequencing library preparation and quantification were performed by Novogene (Sacramento, CA). Illumina NovaSeq 6000 PE150 platform was used to generate paired-end reads. After sequencing, raw data underwent QC to remove low-quality reads, reads with excessive “N”, and adapters to produce high-quality reads, which were then aligned to the *Homo sapiens* reference genome (GRCh38/hg38) using HISAT2 software (version: 2.2.1) for accurate read mapping with the transcriptome. Gene expression levels were normalized as FPKM (fragments per kilobase of transcript sequence per million mapped reads) values to determine the transcript abundance in each sample. Differentially expressed genes (DEGs) were identified using thresholds of fold change ≥ 1.5, log2 fold change ≤ −0.58 or ≥ 0.58, and p-value < 0.05. Further analysis was performed using R programming language within R software (version: RStudio 2025.09.1+401). Visualization of differential expression data through volcano plots and verification of sample grouping results through PCA (Principal component analysis) were performed using ggplot2 (version: 4.0.1). Heat maps were constructed using the pheatmap package. The common overlapping DEGs were identified through the Venn diagram using VennDiagram package. Pathway enrichment analysis using KEGG (Kyoto encyclopedia of genes and genomes) was conducted using Bioconductor packages.

### Antibodies

Primary antibodies used in this study were: anti-STN1 (OBFC1) (cat# WH0079991M1, WB: 1:1000, IF: 1:1000, Sigma-Aldrich), anti-BRCA2 (cat# OP95, WB: 1:10000, EMD Millipore), anti-α-Tubulin (cat# T6199, WB: 1:10000, Sigma Aldrich), anti-Vinculin (cat# 66305, WB, 1:10000, Proteintech), anti-γH2AX-S139 (cat# 2577S, IF: 1:1000, WB: 1:1000, Cell Signaling Technology), anti-myc (cat# 05-419, IF: 1:400, EMD Millipore), pRPA32-S33 (cat# A300-246A, IF:1:4000, WB: 1:10000, Bethyl), Anti-RAD51 (cat# Ab63801, SIRF: 1:200, Abcam), anti-MRE11 (cat# Ab214, SIRF: 1:200, Abcam), anti-CldU (cat# Ab6326, DNA fiber: 1:500, Abcam), anti-IdU (cat# 347580, DNA fiber: 1:50, BD Bioscience), anti-biotin rabbit (cat# 5597S, SIRF: 1:200, Cell Signaling Technology), anti-biotin mouse (cat# SAB4200680, SIRF: 1:200, Sigma-Aldrich). Secondary antibodies used were: Goat anti-Rabbit IgG (H+L) DyLight™ 649 (cat# 35565, IF:1:500, Thermo Scientific), Goat anti-Rabbit IgG (H+L) DyLight™ 550 (cat# 84541, IF: 1:400, Invitrogen), Goat anti-Mouse IgG (H+L) DyLight™ 488 (cat# 35502, IF: 1:800, Thermo Scientific), Alexa Fluor^™^ 568 goat anti-mouse IgG (cat# A-11031, DNA fiber: 1:500, Life Technologies), Alexa Fluor^™^ 588 goat anti-rat IgG (cat# A-11006, DNA fiber: 1:500, Life Technologies), HRP goat anti-mouse IgG (cat# PI-2000, WB: 1:10000, Vector Laboratories), HRP goat anti-rabbit IgG (cat# PI-1000, WB: 1:15000, Vector Laboratories).

### Chemicals

Chemical and drugs used were: Olaparib (cat# SML3705, Sigma-Aldrich), Hydroxyurea (HU) (cat# CAY23725, Cayman Chemicals), CldU (cat# 18155, Cayman Chemicals), IdU (cat# 17125, Sigma-Alderich), EdU (cat# C10337, Invitrogen), Biotin-azide (cat# 1265-5, Bioconjugate Technologies), dimethyl sulfoxide (DMSO) (cat# BP231-1, Fisher Bioreagents^TM^).

### Plasmids and small interfering RNA (siRNA) transfection

WT pCI-neo-Myc-STN1 plasmid was described previously (44). All plasmids were transfected with polyethyleneimine (PEI) with 1:3 DNA:PEI ratio. Knockdown of the target protein using siRNA was carried out using Lipofectamine™ RNAiMAX Transfection Reagent (cat# 13778150, Invitrogen) per manufacturer’s instructions. Briefly, cells were seeded in 6-well plate and incubated overnight to reach ∼60% confluency during the transfection. RNAiMAX and siRNA were separately diluted in Opti-MEM^®^ Reduced-Serum Medium (cat# 31985-061, Gibco). Diluted siRNA and RNAiMAX solutions were mixed after 5 min and incubated further for 15 min at RT. For STN1 knockdown, cells were transfected with siSTN1 (GCTTAACCTCACAACTTAA) once and processed for further experiments after 48 h post transfection. For BRCA2 knockdown, cells were transfected twice with 40 nM siBRCA2 (CAGGACACAATTACAACTAAA, cat# SI02653434, Qiagen) at 24 h intervals and processed for further experiments at 48 h after the second transfection. AllStars Negative Control siRNA (cat# SI103650318, Qiagen) was used as control.

### Western blot analysis

Cells were washed with phosphate-buffered saline (PBS) before being lysed in 1% CHAPS (cat# 808196, MP Biochemicals) buffer (50 mM Tris-HCl, pH 7.4; 110 mM NaCl; 5 mM EDTA; and 1% CHAPS) containing protease inhibitor cocktail (cat# A32963, Thermo Scientific). Whole-cell protein lysates mixed with 2 × SDS loading buffer containing 200 mM Dithiothreitol (DTT, cat# D0632, Sigma-Aldrich) were boiled on 95 °C for 5 min and separated using sodium dodecyl sulfate-polyacrylamide gel electrophoresis (SDS-PAGE) and transferred onto polyvinylidene difluoride (PVDF) membranes (cat# ASEQ83R, Millipore) using ThermoFisher Mini Blot Module (cat# B1000). The membranes were blocked with 5% non-fat dry milk in Tris-buffered saline containing 0.05% Tween 20 (TBST) for 1 h at room temperature. For detection of BRCA2 protein expression, whole-cell lysates were boiled at 55 °C for 10 min in a water bath and then separated on 4-15% gradient gel (cat# 4561083, BioRad). The proteins were then transferred to PVDF membrane at 5 V overnight in transfer buffer containing 10% methanol followed by blocking with 5% non-fat dry milk in TBST for 1 h. The blots were incubated with primary antibodies in TBST overnight at 4 °C. Afterwards, blots were washed with TBST thrice and then incubated with horseradish peroxidase (HRP)-conjugated secondary antibodies for 1 h in 5% non-fat dry milk in TBST at room temperature. The protein bands were then visualized with SuperSignal West Femto Maximum Sensitivity Substrate (Cat#34096, Thermo Scientific). The images were taken by iBright^TM^ imaging system (Invitrogen).

### DNA fiber analysis and S1 nuclease treatment

DNA fiber assays were performed as described previously (51). Briefly, transfected cells expressing the indicated DNA plasmid or siRNA grown in 6-well plate at ∼70% confluency. Cells were labelled with CldU (50 μM) for 20 min followed by washing and subsequent incubation with IdU (250 μM) for 20 min in the culture media. After labelling, cells were immediately washed twice and incubated further with culture media containing 4 mM HU for 3 h. Subsequently, cells were washed once and harvested by trypsinization. Cell pellets were stored in −80 °C until further use.

For S1 nuclease treatment, cells were labelled with CldU (50 μM) for 30 min followed by washing and subsequent incubation with IdU (250 μM) along with HU (0.2 mM for HeLa and 0.5 mM for PEO1) for 150 min. Cells were then washed once with PBS and incubated for 10 min with CSK100 buffer (100 mM NaCl, 3 mM MgCl2, 300 mM sucrose, 10 mM MOPS pH 7.0, 0.5% Triton X-100). Afterwards, cells were washed once again with PBS and incubated for 30 min at 37 °C with S1 nuclease buffer (30 mM sodium acetate, 50 mM NaCl, 10 mM zinc sulphate, 5% glycerol) containing S1 nuclease (cat# 18001016, 50 U/ml, Invitrogen). Cells were harvested by scrapping in PBS containing 0.1% BSA and pelleted for further processing. For DNA fiber stretching, 2 μl of cell suspension (1000 cells/μl in PBS) were spotted on a silanized glass slides and lysed with lysis buffer (200 mM Tris-HCl pH7.4, 0.5% SDS, 50 mM EDTA). DNA fibers were allowed to stretch along the slide surface by tilting at a ∼15° angle and then air-dried. The stretched DNA was fixed with methanol: acetic acid (3:1) for 10 min and denatured in 2.5 M HCl for 80 min. Slides were then washed thrice with PBS and blocked with 5% (w/v) BSA in PBS for 1 h. Nascent DNA were immunostained with anti-CldU and anti-IdU antibodies, and then incubated with anti-rat Alexa Fluor 488 and anti-mouse Alexa Fluor 568 secondary antibodies. Coverslips were mounted with VECTASHIELD Antifade Mounting Medium without DAPI (cat# H-1000, Vector Laboratories). DNA fiber images were taken using a Zeiss Axio Imager M2 epi-fluorescence microscope with a 63 × oil-immersion objective.

### SIRF assay

SIRF assay was performed as previously described (52). Briefly, cells transfected with indicated plasmids or siRNA were grown exponentially on chamber slides and labeled with EdU for 8 min. The cells were washed once with PBS and pre-permeabilized with 0.25% Triton X-100 in PBS for 2 min at room temperature and fixed with 2% paraformaldehyde (PFA, cat# 158127, Sigma-Alderich) for 15 min at room temperature. To induce replication fork stalling, EdU was removed by washing twice with PBS followed by treatment with 4 mM HU for 3 h before fixation. Slides were then washed 3 times in Coplin jars with PBS for 5 min each. After permeabilization with 0.25% Triton X-100 in PBS for 15 min at room temperature, slides were washed 3 times for 5 min each with PBS and then incubated with the click reaction cocktail (2 mM copper sulfate, 10 µM biotin-azide, and 100 mM sodium ascorbate) in a humid chamber at 37 °C for 1 h. Slides were then washed 3 times for 5 min each with PBS and blocked with blocking buffer (10% BSA and 0.1% Triton X-100 in PBS) at 37 °C for 1 h. Primary antibodies were diluted in blocking buffer, dispensed onto slides, and incubated overnight at 4 °C in humidified chamber. For EdU-EdU SIRF, primary antibodies were anti-biotin mouse and anti-biotin rabbit. Slides were then washed three times for 5 min each with wash buffer A (0.01 M Tris, 0.15 M NaCl and 0.05% Tween-20, pH 7.4) and incubated with Duolink in situ PLA^®^ probe anti-mouse MINUS (cat# DUO92002, Sigma-Aldrich) and anti-rabbit PLUS (cat# DUO92002 Sigma-Aldrich) for 1 h at 37 °C. After washing three times with wash buffer A for 5 min each, slides were incubated with Duolink ligation mix (cat# DUO92008, Sigma-Alderich) at 37 °C for 30 min, washed again with wash buffer A two times for 2 min each, incubated with Duolink amplification mix (cat# DUO92008, Sigma-Aldrich) at 37 °C for 100 min, washed with wash buffer B (0.2 M Tris and 0.1 M NaCl) three times for 10 min each and one time in 0.01 × diluted wash Buffer B for 1 min. Lastly, slides were air-dried in the dark and nuclei were counterstaining with VECTASHIELD Antifade Mounting Medium containing DAPI (cat# H-2000, Vector Laboratories). Images were obtained under Zeiss Axio Imager M2 epi-fluorescence microscope with a 63 × oil-immersion objective. SIRF signals from each experiment were normalized to the corresponding EdU-EdU SIRF signals within each replicate to control for background and experimental variation. Normalized values were obtained by expressing the target SIRF signal relative to the EdU-EdU signal from the same assay.

### Immunofluorescence (IF) staining

HeLa cells grown on chamber slides were transfected with indicated plasmids or siRNAs. Cells were fixed with 4% PFA for 15 min and then permeabilized with 0.15% Triton X-100 for 15 min. After washing three times 5 min each with PBS, fixed cells were blocked with 10% BSA for 1 h at 37 °C in a humidified chamber, then incubated with primary antibodies overnight at 4 °C. Samples were washed three times with PBS for 5 min each, followed by incubation with secondary antibodies at room temperature for 1 h. Nuclei were counterstained VECTASHIELD Antifade Mounting Medium with DAPI (cat# H-2000, Vector Laboratories). Fluorescence images were obtained using Zeiss Axio Imager M2 epi-fluorescence microscope with a 63 × oil-immersion objective.

### Clonogenic cell survival assay

HeLa (100 cells/well), PEO1 (150 cells/well), PEO1-R (150 cells/well), and MDA-MB-231 (250 cells/well) were seeded in a 6-well plate 24 h prior to being treated with indicated concentrations of the Olaparib. Cell colonies were allowed to grow for 14 days or until colony size reached a size big enough to stain and count. The colonies were fixed and stained with a solution containing 0.5% crystal violet (cat# C3886, Sigma), 25% Methanol, and double distilled H2O and the number of colonies were counted after washing the stain.

## Results

### RNA-seq analysis of Olaparib-resistant BRCA2-deficient cells reveals distinct and overlapping transcriptional changes in resistant cells

To uncover novel mechanisms underlying PARPi resistance in BRCA2-deficient cells, we sought to take a screening approach by analyzing differentially expressed genes (DEGs) in PARPi-resistant versus sensitive cells, with the goal of identifying candidate genes and pathways that contribute to PARPi resistance. The PEO1 and PEO4 cell lines are a pair of high-grade serous ovarian cancer cell lines derived from the same patient at different stages of chemotherapeutic treatment. While PEO1 is BRCA2-deficient due to a germline nonsense mutation and sensitive to PARPi, PEO4 is BRCA2-proficient and resistant to PARPi due to a revertant mutation in BRCA2. We also developed a new Olaparib-resistant cells (PEO1-R) by chronically exposing PEO1 cells to gradually increasing concentrations of Olaparib over a period of over 6 months, as described in Materials and Methods. Cell proliferation assay showed a significant increase in the IC50 of PEO1-R cells (2.198 µM) compared to the parental PEO1 cells (0.1124 µM), validating its PARPi resistance (Fig. 1A).

**Figure 1.**
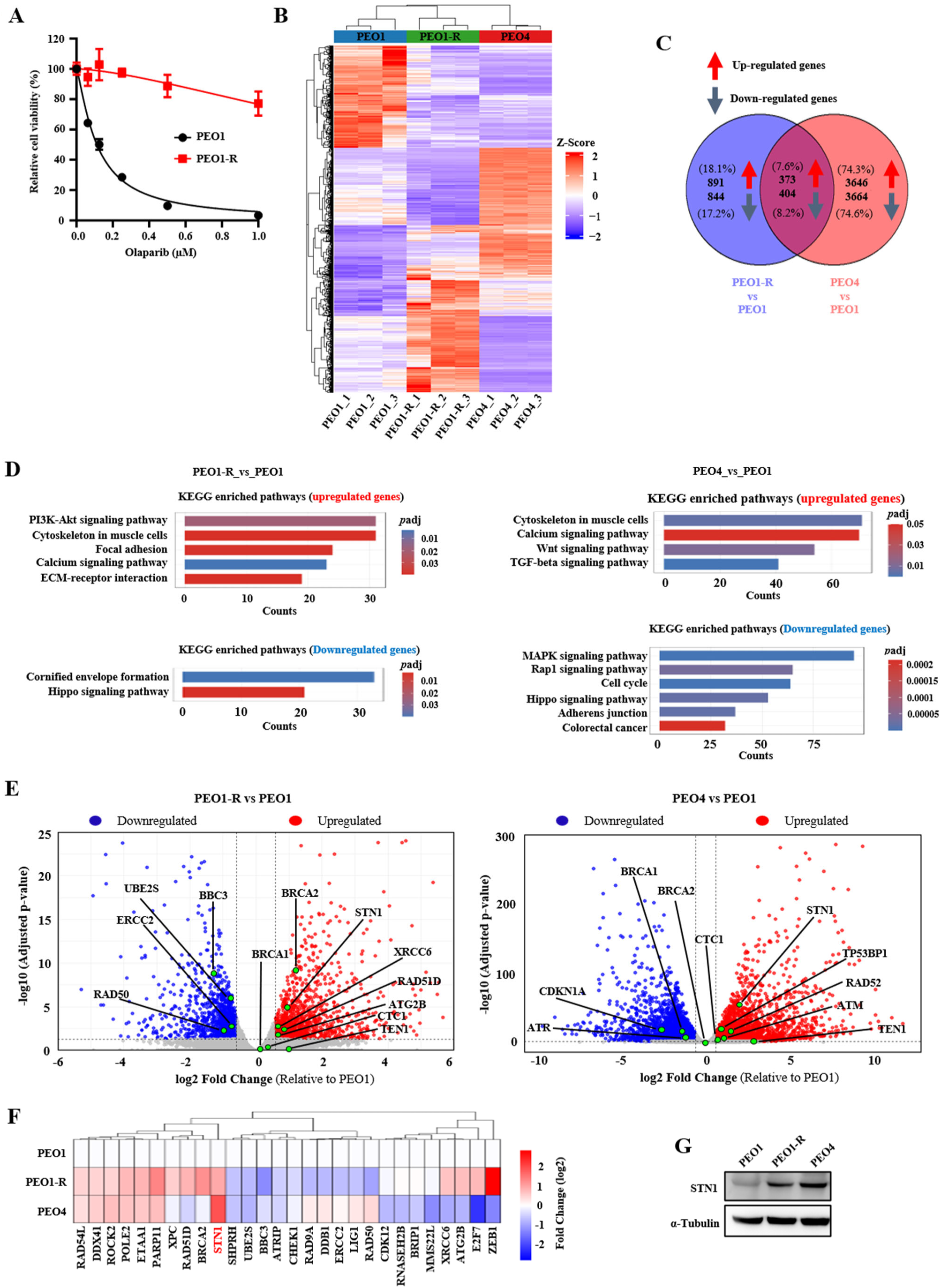
Transcriptomic profiling of PARPi-resistant BRCA2-deficient cells from RNA-seq. **(A)** PEO1 and PEO1-R cell viability was determined after treating cells with various concentrations of Olaparib for 7 days. (**B**) Heat map showing transcriptomic profiles of PEO1, PEO1-R, and PEO4. Color gradient scale represents the log2 fold change. RNA-seq data were generated from three independent biological replicates for each cell line. Statistical significance was set at *p* < 0.05 with a log2 fold change ≥ 0.585. **(C)** Venn diagram illustrating overlapping and unique DEGs between PEO1-R and PEO4 relative to PEO1. (**D)** KEGG pathway enrichment analysis of top significantly enriched pathways among upregulated and downregulated DEGs in PEO1-R and PEO4 when compared to PEO1 cells. Upper panels show the top enriched KEGG pathways for upregulated DEGs, while lower panels show those for downregulated DEGs. X-axis represents gene counts, and y-axis lists pathway names (ordered by significance). Bar color gradient indicates adjusted *p*-value (*p*adj). **(E)** Volcano plot displaying DEGs in PEO1-R and PEO4 relative to PEO1 at the basal level obtained from RNA-seq analysis. Red dots represent significantly upregulated genes; blue dots indicate significantly downregulated genes, and green dots highlight selected genes of interest. (**F)** Heat map showing expressing levels of DEGs involved in genome maintenance and cell survival pathways in PEO1-R and PEO4 cells relative to PEO1. The color gradient scale represents the log2 fold change. (**G)** Western blot analysis showing elevated STN1 protein expression in PEO1-R and PEO4 compared to PEO1.

We then performed RNA-seq using these three isogenic cells, PEO1, PEO4, and PEO1-R. For each cell line, three independent biological replicates were analyzed. Through this analysis, we found that PEO1-R and PEO4, although both resistant to PARPi, showed distinct transcriptome profiles (Fig. 1B and Supplementary Fig. S1). We identified 2,512 DEGs in PEO1-R and 8,087 DEGs in PEO4 compared to the parental PEO1. Venn diagram analysis revealed substantial overlap of DEGs (777 genes) between PEO1-R and PEO4 (Fig. 1C). The distinct transcriptome and DEG profiles in PEO1-R and PEO4 cells suggest that PARPi resistance could arise through more than one molecular route rather than a single, uniform change. It is also possible that the cellular adaptation mechanisms driving resistance may differ between the chronically selected cells (PEO1-R) and cells isolated from the patient following repeated platinum treatments (PEO4).

Although the overall transcriptome and DEG profiles are different, PEO1-R and PEO4 shared limited but notable overlap in DEGs and enriched pathways (Figs. 1C, 1D). KEGG pathway analysis revealed the upregulation of pathways involved in extracellular matrix receptor interaction, focal adhesion, PI3K-Akt signaling, calcium signaling, and cytoskeleton regulation, all of which promote cellular survival (Fig. 1D). In addition, activation of the PI3K-Akt pathway enhances DNA repair capacity and chemoresistance (53–56), while alterations in cytoskeletal dynamics and focal adhesion signaling facilitate resistance-associated cellular adaptation (57–59). In contrast, the tumor-suppressive Hippo signaling pathway was downregulated (Fig. 1D). Suppression of the Hippo signaling pathway induces the nuclear translocation of the transcriptional coactivator yes-associated protein 1 (YAP1), which has been reported to promote cell proliferation and anti-apoptosis, facilitating the development of PARPi resistance in BRCA1/2-mutated cancers (60). These pathway-level changes complement the genetic alterations, collectively enabling cancer cells to withstand Olaparib treatment.

### Distinct and overlapping changes in DDR and DNA repair genes in PEO1-R and PEO4

Next, we analyzed DEGs and pathway changes in DDR and DNA repair in the two resistant cell lines. Volcano plots revealed distinct DEG profiles in PEO1-R and PEO4 compared with PEO1 (Fig. 1E and Supplementary Fig. S2). Notably, BRCA1 expression levels remained unchanged in both cell lines (Fig. 1E). In addition, PEO1-R cells showed a unique upregulation of DEGs across multiple DNA repair pathways including HR (RAD51D), NHEJ (XRCC6/Ku70), nucleotide excision repair (NER) (XPC), mismatch repair (MLH1), and cell survival and autophagy pathways (ATG2B, HPF1). In contrast, the upregulated DEGs unique to PEO4 cells were involved in HR (RAD52), DNA damage sensing and checkpoint activation (ATM, TP53BP1), NHEJ (XRCC5/Ku80), base excision repair (BER) (APEX1, LIG3), NER (ERCC4), MMR (PMS2), cell cycle regulation (TP53) and DDR signaling and chromatin remodeling (KAT5, H2AFX) (Supplementary Fig. S2). These results indicate that the development of drug resistance does not rely on a single pathway but instead can involve distinct pathways. Furthermore, the coexistence of both shared and distinct transcriptional changes in the two cell lines suggests that their responses to different chemotherapeutic agents may be driven by overlapping yet agent-specific molecular mechanisms.

### STN1 is upregulated in Olaparib-resistant BRCA2-deficient cells

We then focused on overlapping DEG candidates shared between PEO1-R and PEO4, particularly those involved in DDR and repair. Significant upregulation was observed in several DDR and repair genes such as RAD54L, ETAA1 and POLE2, as well as in cell survival related gene like ROCK2 (Fig. 1F). Interestingly, STN1, but not other CST components (CTC1 and TEN1), was consistently upregulated in both PEO1-R and PEO4 cells (Figs. 1E, 1F). Western blot analysis confirmed significantly upregulated STN1 protein levels in Olaparib-resistant PEO1-R and PEO4 cells, validating the transcriptomic findings (Fig. 1G). Due to the unavailability of reliable CTC1 and TEN1 antibodies, we were unable to directly assess CTC1 and TEN1 protein levels in these cell lines.

### Ectopic overexpression of STN1 correlates with Olaparib resistance and BRCA2 restoration

As STN1 was upregulated in Olaparib-resistant PEO1-R and PEO4 cells, we sought to determine whether STN1 overexpression contributes to the development of PARPi resistance. Using retroviral transduction, we stably overexpressed STN1 in PEO1 cells to the levels comparable to those in PEO1-R and PEO4 (Fig. 2A). Colony formation assay showed that ectopic overexpression of STN1 in PEO1 cells significantly increased Olaparib resistance, compared to control cells (Figs. 2B-C), suggesting that increasing STN1 abundance may lead to PARPi resistance in BRCA2-deficient cells.

**Figure 2.**
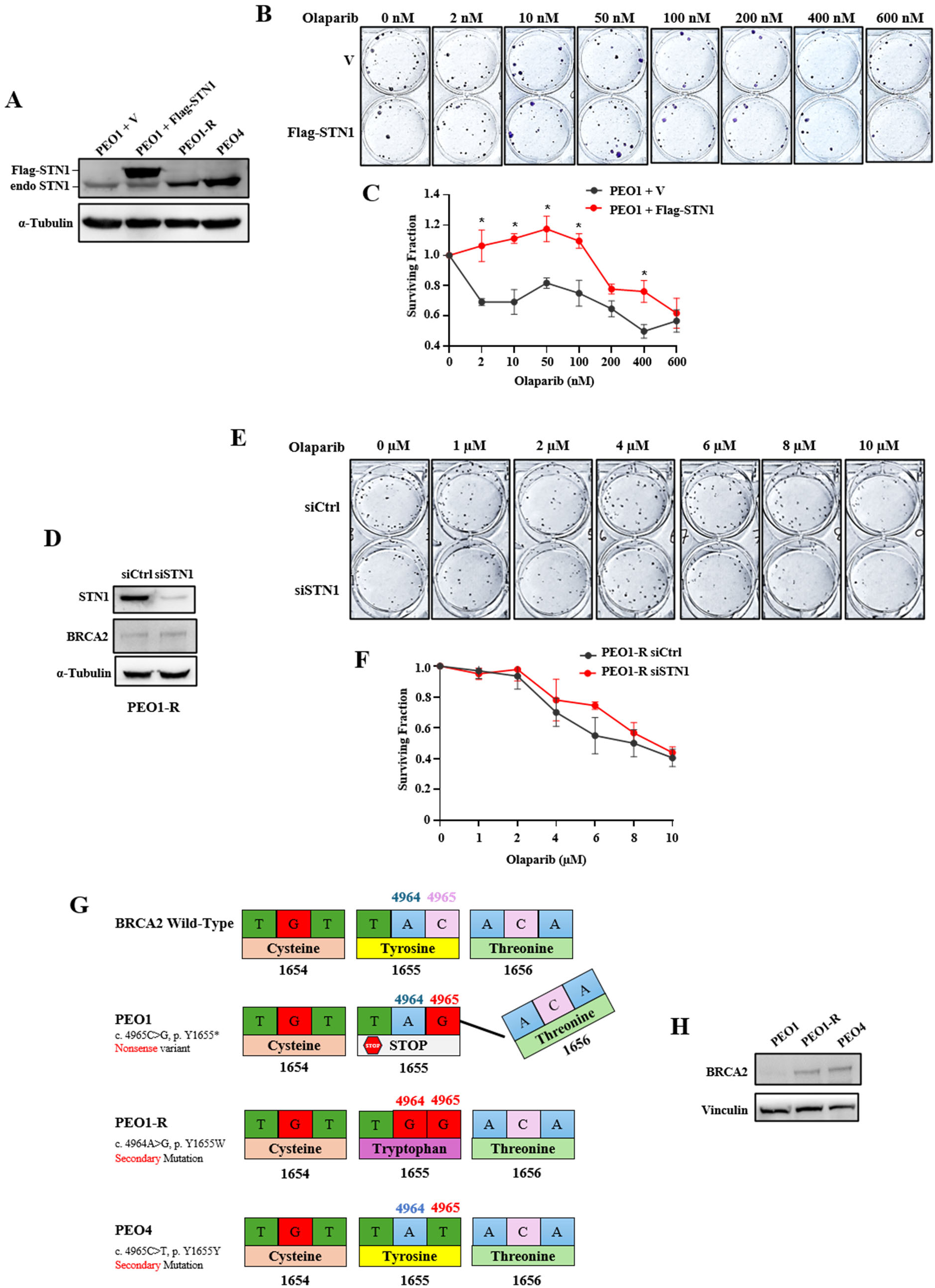
Upregulation of STN1 correlates with Olaparib resistance and BRCA2 restoration. (**A**) Western blot shows that STN1 levels in PEO1 cells stably expressing exogenous Flag-STN1 are comparable to the elevated STN1 levels in PEO1-R and PEO4 cells. Endo-STN1: endogenous STN1. (**B**) Representative images of clonogenic assay of PEO1 expressing vector or overexpressed Flag-STN1 after treatment with indicated concentrations of Olaparib. Quantification of survival is shown in **(C)**. In all survival curves, data are mean ± SEM of at least two independent experiments. One-way ANOVA was used to determine the significance. (^∗^*P* < 0.05). (**D**) Western blot shows the STN1 knockdown and BRCA2 expression in PEO1-R cells. (**E**) Representative images of clonogenic assay of PEO1-R transfected with control siRNA and siSTN1 and treated with indicated concentrations of Olaparib. Quantification of survival after STN1 depletion is shown in **(F)**. (**G**) Illustration of BRCA2 sequencing results showing BRCA2 reversion mutations in PEO1-R and PEO4 cells. (**H**) Western blot shows BRCA2 restoration in PEO1-R and PEO4 cells.

Next, we checked whether downregulation of STN1 in PEO1-R cells could restore sensitivity to Olaparib. As shown in Figs. 2D-F, STN1 depletion in PEO1-R cells did not re-sensitize cells to Olaparib. This result encouraged us to explore the underlying mechanism driving Olaparib resistance in these cells. Western blot analysis revealed that PEO1-R cells expressed full length BRCA2 (Fig. 2D), indicative of BRCA2 restoration. Depletion of STN1 in PEO1-R cells had no effect on BRCA2 expression levels (Fig. 2D). Previous studies have shown that the parental PEO1 cells carry the Y1655X (c.4965C > G) mutation in *BRCA2*, resulting in a premature stop codon and leading to a truncated protein. On the other hand, PEO4 shows a secondary mutation at c.4965C>T which results in a TAT codon encoding tyrosine (Y), indicating restoration of full-length WT BRCA2 (28,29) (Fig. 2G). Our sequencing results of BRCA2 in PEO1-R showed the secondary mutation at c.4964A>G, which resulted in a TGG codon translating into tryptophan (W), thereby yielding a full-length BRCA2 (Figs. 2G, 2H). A similar *BRCA2* reversion mutation (c.4964A > T, leading to Y1655F), which restored full-length BRCA2, has previously been reported after long-term Olaparib treatment (29). The restoration of functional BRCA2 in PEO1-R cells probably allowed cells to fully regain HRR proficiency and fork stability, so that depleting STN1 alone was not able to significantly affect PARPi sensitivity.

Next, we tested whether PAPRi resistance is specific to STN1 or extends to other components of the CST complex. Ectopically expressing CTC1 in PEO1 cells, achieved either by retroviral transduction or transient transfection, did not confer Olaparib resistance (Supplementary Fig. S3). However, due to the unavailability of reliable CTC1 antibodies, we were not able to verify CTC1 overexpression (Supplementary Fig. S4). In addition, it has been reported that STN1 protects CTC1 from degradation by the TRIM32 ubiquitin E3 ligase (61). Therefore, it is possible that ectopically expressed CTC1 was unstable with limited STN1 in cells, limiting its ability to promote PARPi resistance.

### STN1 overexpression increases Olaparib resistance in multiple BRCA2-deficient cancer cell lines

We next sought to determine whether Olaparib resistance driven by STN1 upregulation was limited to specific cell types or represented a broader mechanism. Ectopic overexpression of STN1 in BRCA2-depleted triple-negative breast cancer cell line MDA-MB-231 led to significantly increased cell survival compared to BRCA2-depleted cells transfected with empty vector (Fig. 3). In addition, ectopic expression of STN1 in BRCA2 KO HeLa cells resulted in a mild increase of Olaparib resistance at low concentrations (1 µM) (Supplementary Fig. S5). Collectively, these findings support that elevated STN1 expression increases Olaparib resistance in various BRCA2-deficient cells.

**Figure 3.**
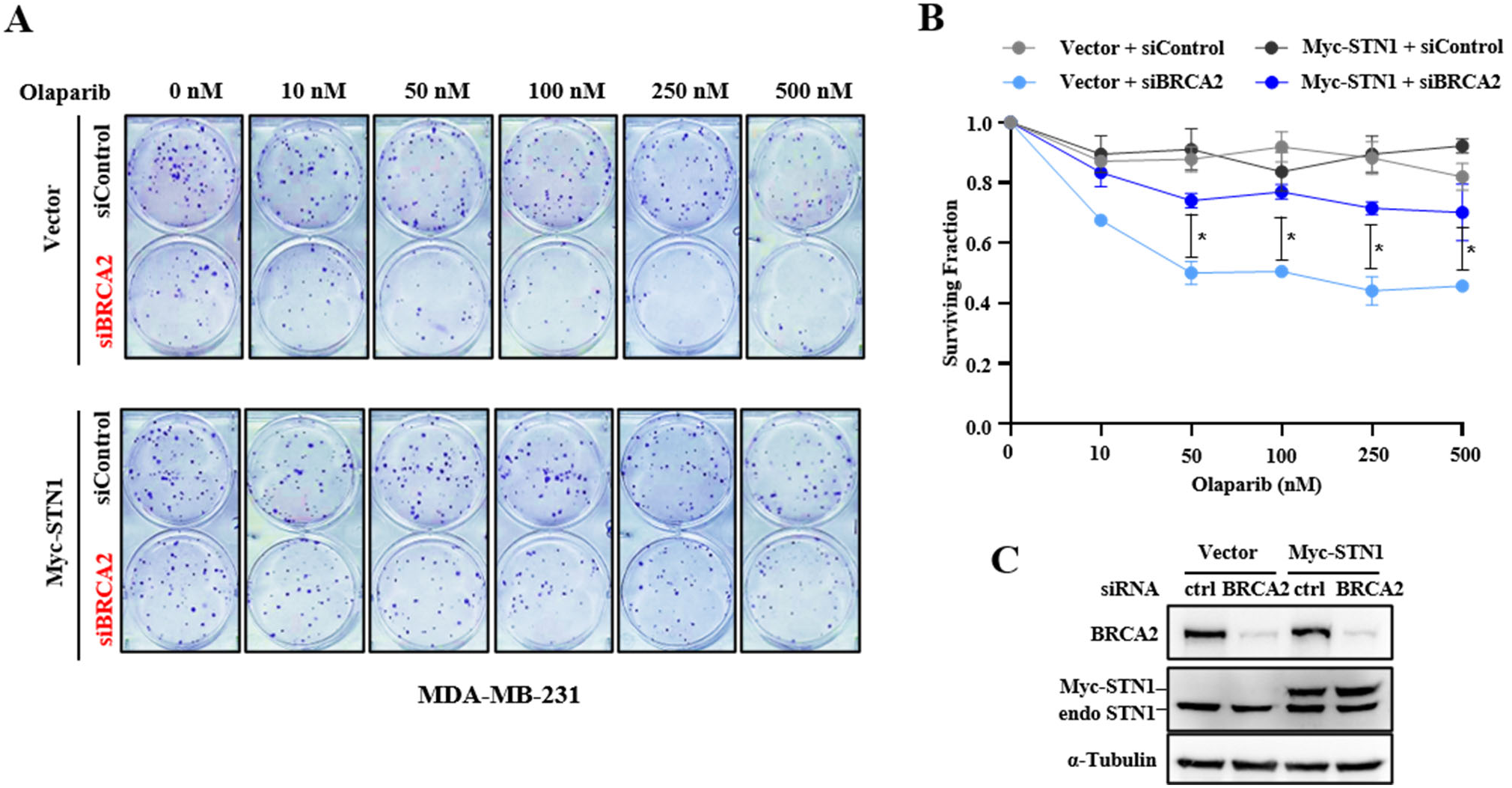
STN1 overexpression increases Olaparib resistance in BRCA2-deficient MDA-MB-231 cells. **(A)** Clonogenic survival of BRCA2 depleted MD-MB-231 cells with or without Myc-STN1 overexpression. Quantification of survival (mean ± SEM from two independent experiments) is shown in **(B)**. *P* values: One-way ANOVA. ^∗^*P* < 0.05. **(C)** Western blot showing STN1 overexpression and BRCA2 knockdown in MD-MB-231 cells

### STN1 overexpression decreases DNA damage levels in BRCA2-deficient cells

PARPi such as Olaparib induce DNA damage by trapping PARP1 and PARP2 at single strand breaks, leading to replication fork collapse and DSB accumulation that cannot be efficiently repaired in BRCA-deficient cells due to impaired HRR (6,62). To examine the role of STN1 overexpression in replication fork stability under replication stress in BRCA2-deficient cells, we measured γH2AX levels in PEO1 cells with and without stable overexpression of STN1 in PEO1 cells. STN1 overexpression showed a drastic reduction of γH2AX levels upon HU treatment (Figs. 4A-C). Similar γH2AX reduction was also observed in BRCA2-depleted HeLa cells with concurrent STN1 overexpression (Figs. 4D-F). These results indicate that increased STN1 abundance in BRCA2-deficient cells reduces DNA damage.

**Figure 4.**
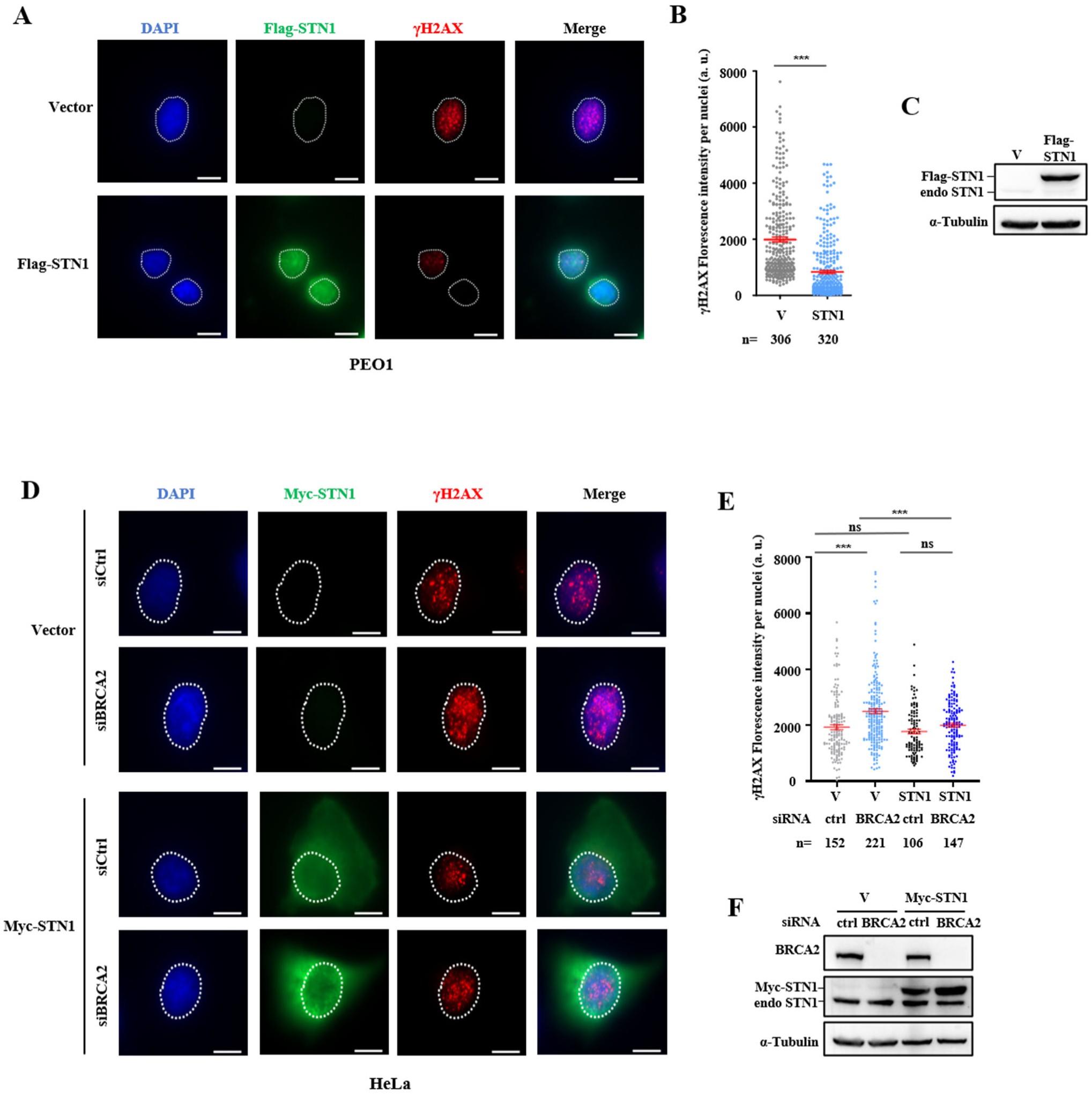
STN1 overexpression reduces γH2AX formation in BRCA2-deficient cells. **(A)** Representative IF images (scale bar, 10 μm) of γH2AX staining upon HU treatment (4 mM, 3 h) in PEO1 cells stably expressing vector or Flag-STN1. Nuclei are outlined with white dash lines. **(B)** Quantification of fluorescence intensity of γH2AX per nuclei. Data presented as mean ± SEM. *n*: number of cells analyzed. *P* values were calculated by Mann-Whitney test. ^∗∗∗^*P* < 0.001. **(C)** Western blot of STN1 confirming STN1 overexpression in PEO1 cells expressing Flag-STN1. **(D)** Representative IF images (scale bar, 10 μm) of γH2AX staining upon HU treatment (4 mM, 3 h) in HeLa cells transfected with vector or Myc-STN1 followed by BRCA2 depletion. Nuclei are outlined with white dash lines. **(E)** Quantification of fluorescence intensity of γH2AX per nuclei. Data presented as mean ± SEM. *n*: number of cells analyzed. *P*: one-way ANOVA with Tukey’s multiple comparisons test. ^∗∗∗^*P* < 0.001. **(F)** Western blot confirming Myc-STN1 overexpression and siRNA-mediated BRCA2 knockdown in HeLa cells.

### STN1 overexpression counteracts fork degradation in BRCA2-deficient cells

BRCA2 is required for RAD51 loading to stalled forks to prevent MRE11-mediated degradation of nascent strand DNA (49,63–65). Similarly, CST facilitates RAD51 loading to stalled forks to antagonize unscheduled MRE11 degradation of nascent strand DNA (43,66). We therefore investigated whether STN1 overexpression could restore fork stability in BRCA2-deficient cells using DNA fiber assays. Our results showed that upon HU treatment, BRCA2-deficient cells exhibited a decreased IdU/CldU ratio, indicative of HU-dependent nascent DNA degradation (Figs. 5A-B). Notably, STN1 overexpression rescued such degradation (Figs. 5A-B). This phenotype was not restricted to specific cell types as we observed the same phenotype in BRCA2-depleted HeLa cells (Fig. 5C). These findings suggest that STN1 overexpression mitigates replication stress-associated fork degradation in BRCA2-deficient cells.

**Figure 5.**
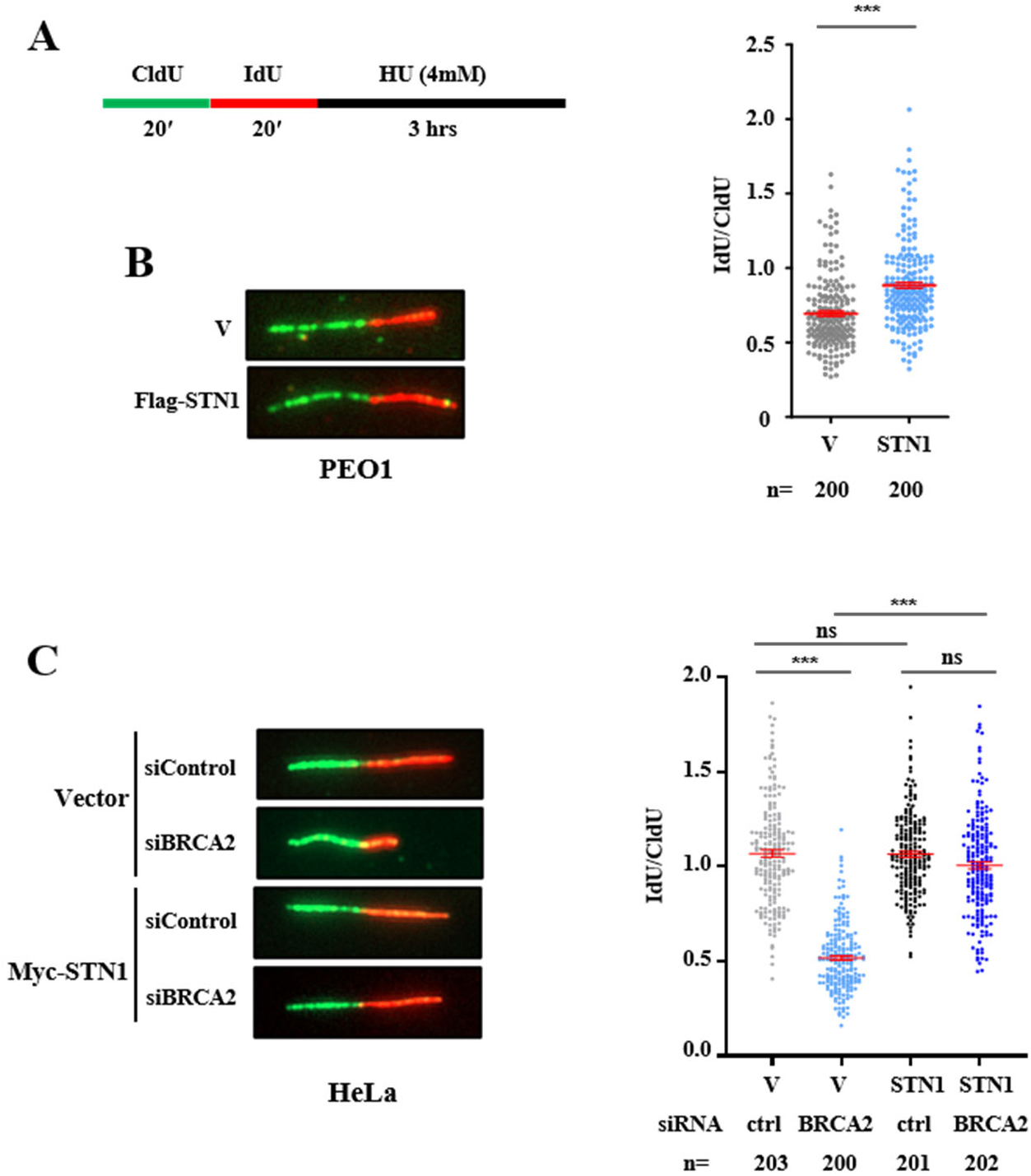
STN1 overexpression counteracts fork degradation in BRCA2-deficient cells.(A) Scheme of DNA fiber assay for analyzing nascent strand DNA degradation. **(B)** Representative DNA fiber images in PEO1 cells. Scatter plots showing the IdU/CldU ratio in PEO1 cells stably expressing vector or Flag-STN1 following HU treatment. The graphs are representative of two independent experiments. *n* represents the number of DNA fibers analyzed in each sample in each experiment. Statistical significance was determined by Mann-Whitney test (^∗∗∗^*P* < 0.001). **(C)** Representative images of DNA fiber in HeLa cells. Scatter plots shows the IdU/CldU ratio in HeLa cells expressing vector or Myc-STN1 with and without BRCA2 depletion following HU treatment. The graphs are representative of two independent experiments. *n* represents the number of DNA fibers analyzed in each sample in each experiment. Statistical significance was determined by one-way ANOVA (^∗∗∗^*P* < 0.001).

### STN1 overexpression partially rescues the localization of RAD51 and MRE11 at stalled forks caused by BRCA2 loss

RAD51 loading is essential for protecting reversed replication forks from MRE11-dependent nucleolytic degradation (21,22). Since BRCA2 promotes the formation of RAD51 filaments on replication forks to prevent MRE11-dependent fork degradation (22,23), we examined the effects of STN1 overexpression on RAD51 and MRE11 association to stalled forks in the absence of BRCA2. RAD51 SIRF analysis revealed that STN1 overexpression in PEO1 cells markedly increased RAD51 SIRF foci, whereas MRE11 association with stalled forks were reduced (Figs. 6A-C). Similar effects were also observed after STN1 overexpression in HeLa cells depleted of BRCA2 (Figs. 6D, 6E). These findings indicate that STN1 overexpression counteracts the effects of BRCA2 loss, at least in part, by enhancing RAD51 recruitment and limiting MRE11-mediated degradation, thereby stabilizing replication forks in BRCA2-deficient cells.

**Fig. 6.**
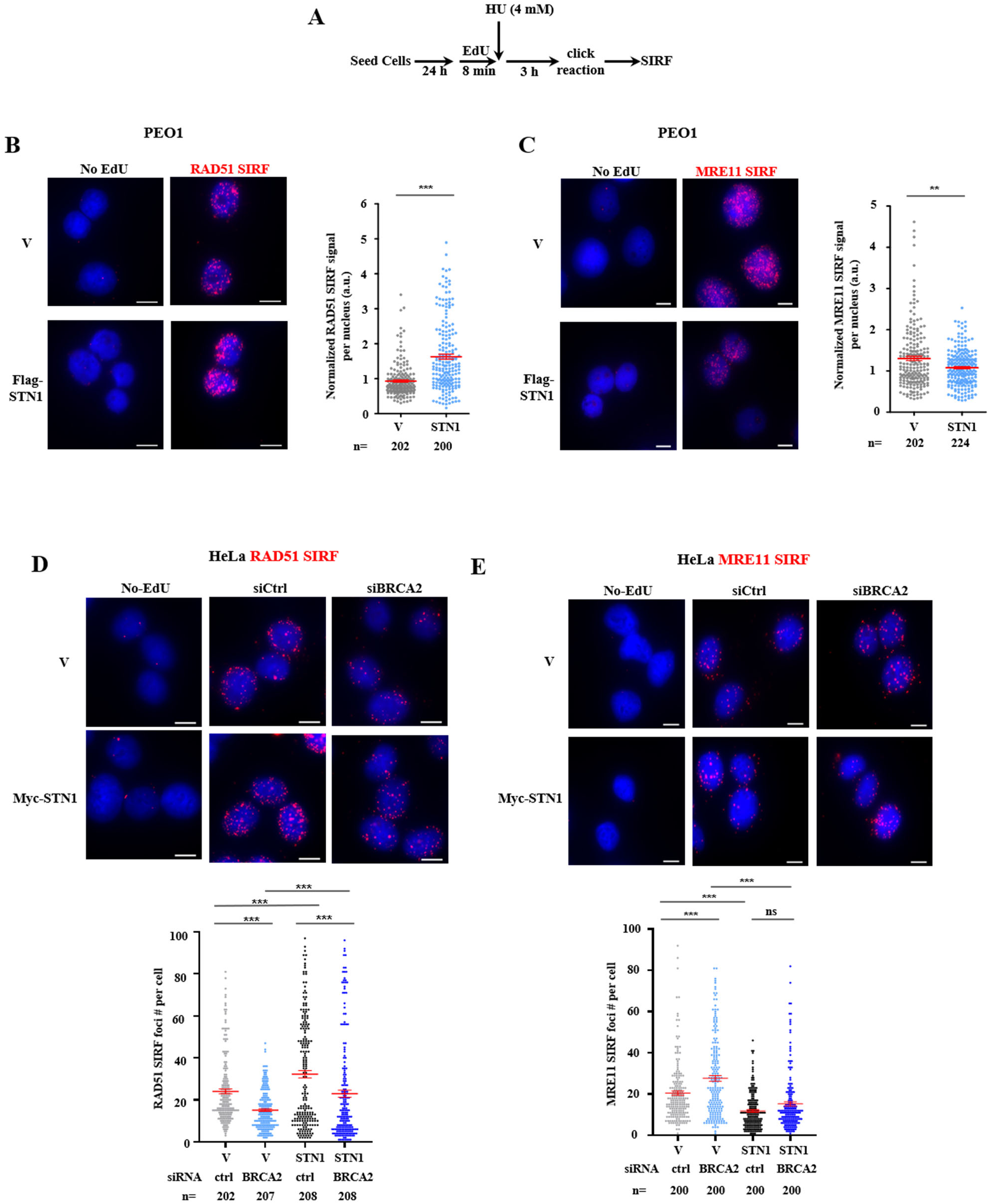
STN1 overexpression partially rescues the localization defects of RAD51 and MRE11 at replication forks caused by BRCA2 loss. **(A)** Scheme of SIRF experiments. **(B)** Representative images of RAD51 SIRF showing RAD51 localization at nascent DNA in BRCA2-deficient PEO1 cells and the rescue effect by stable overexpression of Flag-STN1. Scale bar: 10 μm. Scatter plot shows RAD51 SIRF signals normalized to EdU-EdU SIRF signals for each nucleus. **(C)** Representative images of MRE11 SIRF showing MRE11 localization at nascent DNA in BRCA2-deficient PEO1 cells and the rescue effect by stable overexpression of Flag-STN1. Scale bar: 10 μm. Scatter plot shows MRE11 SIRF signals normalized to EdU-EdU SIRF signals for each nucleus. Mean ± SEM values for each sample are shown in red. *n* represents the number of cells analyzed in each sample. *P* values were calculated by Mann-Whitney test. \*\*\**P* < 0.001. **(D)** Representative images of RAD51 SIRF showing RAD51 localization at nascent DNA in BRCA2-deficient HeLa cells and the rescue effect by STN1 overexpression. Scale bar: 10 μm. Scatter plot shows RAD51 SIRF signals normalized to EdU-EdU SIRF signals for each nucleus. **(E)** Representative images of MRE11 SIRF showing MRE11 localization at nascent DNA in BRCA2-deficient HeLa cells and the rescue effect by STN1 overexpression. Scale bar: 10 μm. Scatter plot shows MRE11 SIRF signals normalized to EdU-EdU SIRF signals for each nucleus. Mean ± SEM values for each sample are shown in red. *n* represents the number of cells analyzed in each sample. *P* values were calculated by Mann-Whitney test. \*\*\**P* < 0.001.

### STN1 overexpression suppresses ssDNA gap accumulation in BRCA2-deficient cells

In addition to its key roles in HR and fork protection, BRCA2 suppresses ssDNA gap formation in the genome (7,8,25,26). SsDNA gaps can arise from stalling of DNA polymerases or uncoupling of leading and lagging strand synthesis during persistent replication stress, Okazaki fragment maturation defects, PrimPol-mediated repriming downstream of DNA lesions, impaired chromatin assembly during replication stress, and others (9). If unrepaired, these ssDNA gaps are further expanded by nucleases, contributing to genomic instability (66–69). BRCA2 suppresses gap formation via multiple mechanisms. First, BRCA2 promotes RAD51 filament formation to protect ssDNA while displacing RPA. BRCA2-RAD51 inhibits nucleolytic degradation of DNA to prevent ssDNA formation. Second, by interacting with MCM10, BRCA2 suppresses PrimPol activity to prevent gap accumulation (70). Third, BRCA2 stimulates replication-associated homology-directed repair. In addition, BRCA2 has been implicated in facilitating POLθ-mediated gap filling by limiting MRE11 extension of gaps into longer stretches of ssDNA (26,71,72). Recent studies have shown that ssDNA gap accumulation in BRCA2-deficient cells plays an important role in PARPi sensitivity, as cells resistant to PARPi often restore ssDNA gap suppression, not necessarily HRR (7,8).

To determine whether STN1 overexpression could alleviate ssDNA gap formation caused by BRCA2 loss, we first analyzed RPA32 Ser33 phosphorylation (pS33) by immunofluorescence analysis, a marker of ssDNA accumulation at replication-associated lesions (73,74), in HU-treated PEO1 cells with or without stable STN1 overexpression, as well as in HeLa cells following BRCA2 depletion, with and without concurrent STN1 overexpression. We observed that overexpressing STN1 in PEO1 significantly reduced pRPA32 fluorescence intensity compared to the control (Fig. 7A). Western blot analysis further confirmed increased pRPA32 levels in HU treated PEO1 cells, and STN1 overexpression reduced pRPA32 (Fig. 7B). Likewise, STN1 overexpression in BRCA2-deficient HeLa cells significantly reduced pRPA32 caused by BRCA2 depletion (Fig. 7C). These results suggest that increased STN1 expression mitigates ssDNA accumulation caused by BRCA2 loss (Fig. 7C).

**Fig. 7.**
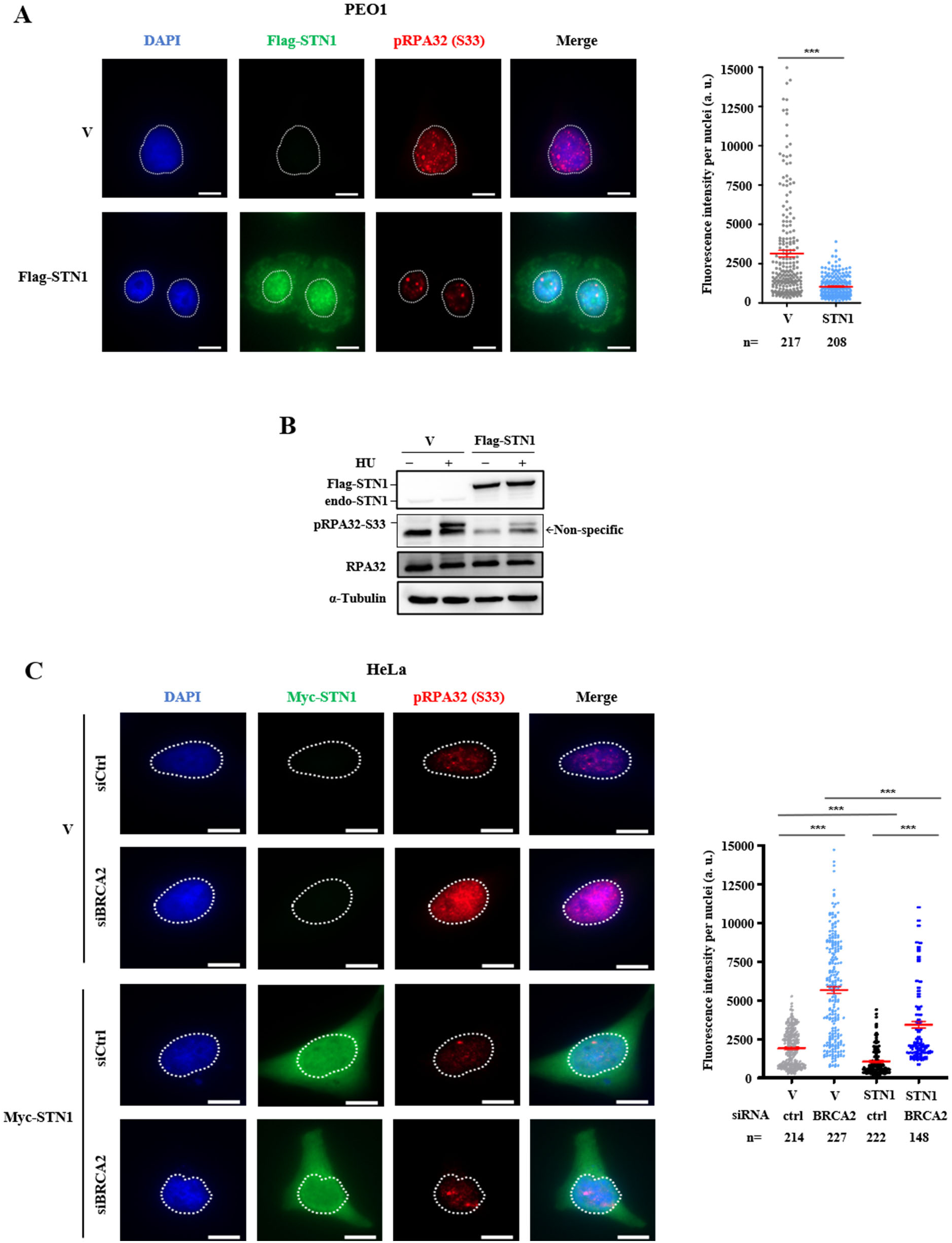
STN1 overexpression counteracts ssDNA generation in BRCA2-deficient cells. **(A)** Representative images from immunofluorescence staining for RPA32 pS33 in PEO1 cells stably expressing vector or Flag-STN1. Cells were treated with HU (4 mM, 3 h) prior to fixation. Nuclei are outlined with white dash lines. Scatter plot shows pS33 RPA32 fluorescence intensity (a. u.) per nucleus. Mean ± SEM values for each sample are shown in red. *n*: the number of cells analyzed in each sample. *P* values were calculated by Mann-Whitney test. \*\*\**P* < 0.001. **(B)** Western blot analysis of RPA32 pS33 upon HU treatment (4 mM, 3 h) in PEO1 cells with stable expression of Flag-STN1. **(C)** Representative images from immunofluorescence staining for RPA32 pS33 in HeLa cells overexpressing Myc-STN1 with or without BRCA2 depletion. Cells were treated with HU (4 mM, 3 h) prior to fixation. Nuclei are outlined with white dash lines. Scatter plot shows pS33 RPA32 fluorescence intensity (a. u.) per nucleus. In Myc-STN1 expressing samples, only cells with green fluorescence (indicating Myc-STN1 overexpression) were included in analysis. Mean ± SEM values for each sample are shown in red. *n*: the number of cells analyzed in each sample. *P* values: one-way ANOVA. \*\*\**P* < 0.001.

To further validate the role of STN1 in reducing ssDNA gaps, we performed S1 nuclease DNA fiber assays, which detect ssDNA gaps formed in replicating DNA due to their sensitivity to S1 nuclease digestion (8). Cells were pulsed labeled with CldU for 30 min, followed by IdU labeling in the presence of HU for an extended period (150 min) to account for slowed replication progression (Fig. 8A). S1 nuclease treatment was performed prior to cell collection for DNA fiber analysis. Consistent with previous reports, reduced IdU tract lengths were observed in PEO1 cells (Figs. 8B-D) and HeLa cells (Fig. 8E-G) following S1 nuclease treatment, indicative of increased ssDNA gap formation. Notably, STN1 overexpression restored S1-sensitive IdU tract lengths in both cell lines (Figs. 8B-8G), indicating a protective effect against ssDNA gap accumulation. Collectively, these findings demonstrate that increasing STN1 expression counteracts ssDNA gap accumulation in BRCA2-deficient cells.

**Fig. 8.**
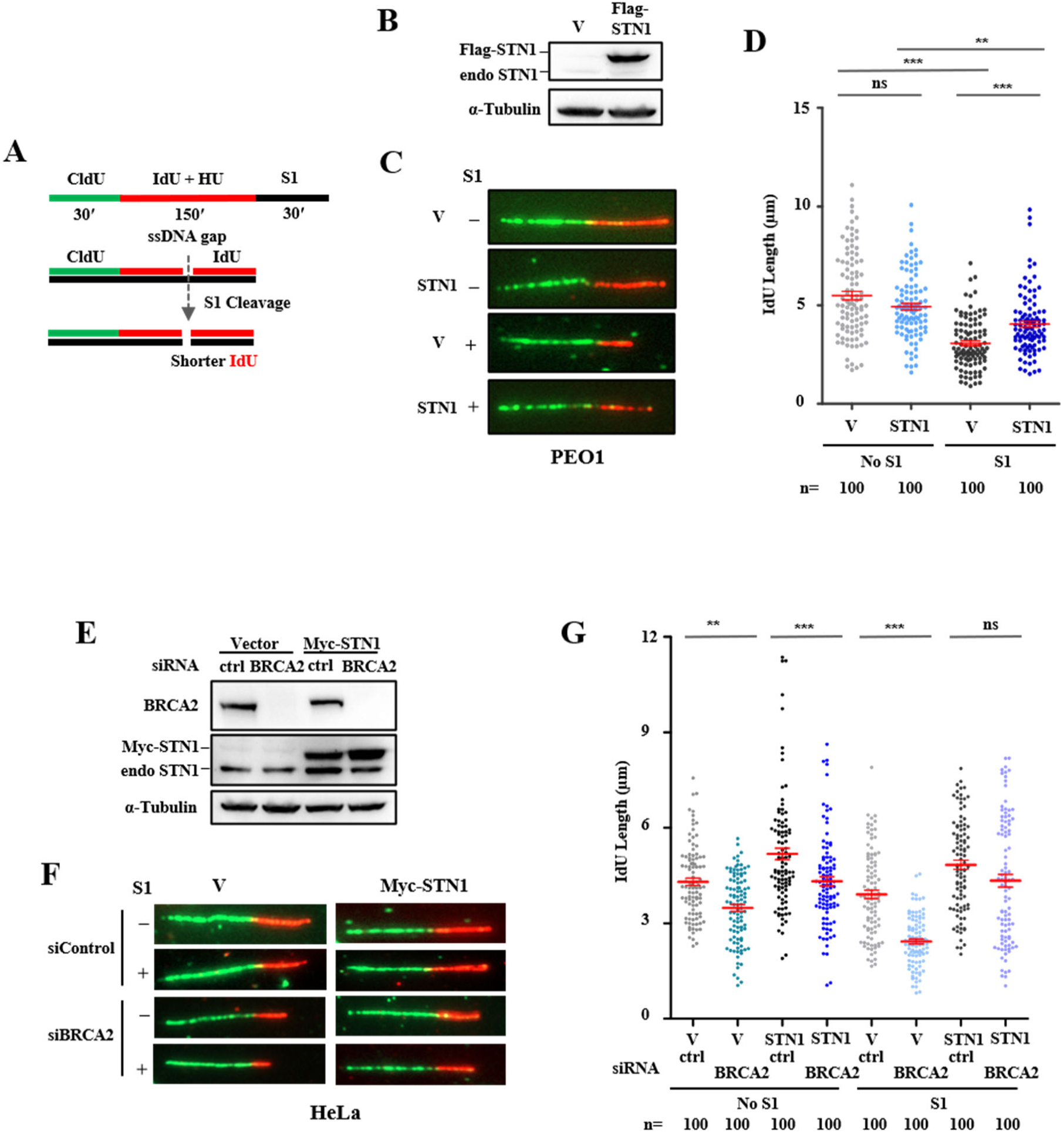
STN1 overexpression counteracts ssDNA gap accumulation in BRCA2-deficient cells. **(A)** Scheme of the CldU-IdU labelling and S1 nuclease treatment strategy for detecting ssDNA gaps upon HU treatment. **(B)** Western blot confirming Flag-STN1 overexpression in PEO1 cells. **(C)** Representative images of S1 DNA fiber in PEO1 cells. (**D)** Scatter plots showing IdU tract lengths in PEO1 cells stably expressing vector or Flag-STN1, treated with HU (0.5 mM), and analyzed with or without S1 nuclease (*n* = 100). **(E)** Western blot analysis confirming the overexpression of Myc-STN1 and BRCA2 depletion in HeLa cells. (**F)** Representative S1 DNA fiber images in HeLa cells. **(G)** Scatter plots showing IdU tract lengths in HeLa cells transfected with indicated plasmids and siRNAs, treated with HU (0.2 mM), and analyzed with or without S1 nuclease (*n* = 100).

## Discussion

Although PARPi selectively target HRR-deficient cancers, resistance frequently emerges, limiting their clinical efficacy. Understanding the molecular mechanisms that enable HR-deficient cells to bypass replication stress and survive PARPi treatment remains a central challenge. In this study, we identify a previously unrecognized mechanism that STN1 overexpression facilitates BRCA2-deficient cells to acquire PARPi resistance. STN1 is a component of the ssDNA binding protein complex CST. Previously we have demonstrated that CST directly interacts with RAD51 and promotes RAD51 recruitment to stalled replication forks to suppress MRE11-mediated nucleolytic degradation (43,75). Using RNA-seq, we identify that STN1 is upregulated in PARPi-resistant BRCA2-deficient ovarian cancer cells. Our work shows that overexpressing STN1 functionally compensates for the loss of BRCA2 by restoring RAD51 loading to stalled forks, suppressing MRE11 access to forks and nascent strand degradation, diminishing the formation of ssDNA gaps, and reducing the accumulation of γH2AX foci, indicating that STN1 overexpression decreases replication stress-induced DNA damage in BRCA2-deficient cells. Taken together, our results support a model that STN1 overexpression partially compensates for the loss of BRCA2, thereby attenuating the cytotoxic consequences of PARP inhibition.

Our finding suggests that STN1 may offer an alternative fork protection pathway by decreasing the dependence on BRCA2 for maintaining fork stability and enabling cells to tolerate PARPi-induced replication perturbations. While BRCA2 plays essential roles in both fork protection and HRR, our findings suggest that bypassing the fork protection defect and/or ssDNA gap accumulation may promote acquired PARPi resistance. This model aligns with emerging evidence that fork protection restoration, for example, via loss of fork remodelers such as SMARCAL1 or ZRANB3, loss of nucleases like PTIP, or increased expression of replication stress response proteins, confers resistance even in HR-deficient tumors (6,23,76).

While our data suggest that STN1 overexpression facilitates PARPi-sensitive BRCA2-deficient cells to acquire resistance, the reverse does not appear to be true, as depletion of STN1 in PARPi-resistant cancer cells that have already restored BRCA2 expression does not re-sensitize these cells to PARPi (Fig. 2E). We speculate that once BRCA2 function has been restored, the BRCA2-RAD51 pathway is sufficient to protect forks, thereby rendering CST dispensable for the maintenance of PARPi resistance. This is consistent with studies showing that PARPi-resistant tumors often acquire multiple, often redundant mechanisms, including BRCA reversion, suppression of 53BP1-Shieldin-mediated end protection, replication fork stabilization, and drug efflux or PARP1 alterations (77). Consequently, disrupting a single compensatory factor is insufficient to reverse resistance once a stable resistant state has been established. It is also possible that STN1 functions primarily during the transitional phase of resistance acquisition, in particular when BRCA2-deficient cells experience severe replication stress, and that once resistant cells have undergone broader transcriptional, epigenetic, or genomic remodeling that re-establishes replication homeostasis, STN1 overexpression is no longer needed. Alternatively, stable PARPi-resistant cells may upregulate additional fork protection factors or alter nucleases such as MRE11 or DNA2 in ways that bypass the need for STN1. These possibilities highlight that STN1 overexpression may act as an initiating or facilitating factor to promote survival during the early stages of resistance development, whereas long-term, fixed resistance states rely more on BRCA2-restorative mechanisms.

It is noteworthy that only STN1, but not the other two components of the CST complex, is sufficient to promote PARPi resistance (Fig. 1, Supplementary Fig. S3). Consistent with this observation, a recent study reports that elevated STN1 expression, but not CTC1 or TEN1, facilities metastasis in pancreatic cancer (78). We think this phenomenon may be explained by the underlying molecular interactions within the CST complex. Within the CST complex, CTC1 and TEN1 do not interact directly. Instead, the overall complex assembly and stability reply on STN1, which acts as a bridging subunit between CTC1 and TEN1 (79). In addition, STN1 is essential for maintaining CTC1 protein stability (61). Consequently, elevated STN1 levels may enhance CST function by increasing CTC1 stability while simultaneously increasing the availability of TEN1 to associate with the CTC1-STN1 subcomplex, thus increasing the overall pool of functional CST complexes capable of binding DNA and stalled replication forks to protect nascent strand DNA from degradation and suppress gap accumulation. However, it is also possible that STN1 may compensate for BRCA2 loss through a CST complex-independent function, as a recent report demonstrates that STN1 participates in DSB repair independently of CTC1 and TEN1 (80). Currently, our current data do not distinguish whether STN1 acts independently of the CST complex or whether increased STN1 expression enhances CST function by increasing the availability of a limiting component. Definitively resolving this question is challenging because the CST complex plays essential roles in DNA replication and genome maintenance. Disruption of individual CST components compromises cell proliferation and replication fitness, making it difficult to separate CST-dependent effects on PARPi response from broader consequences of CST dysfunction. In addition, the lack of reliable CTC1 and TEN1 antibodies makes it difficult to directly evaluate the roles of individual CST components in mediating PARPi response. Therefore, while our data identify STN1 as the only CST component whose upregulation is associated with and sufficient to promote PARPi resistance-associated phenotypes, additional studies will be required to determine whether these effects reflect a unique function of STN1 or enhanced activity of the CST complex as a whole.

Our findings in BRCA2-deficient cells are a stark contrast to the phenotype observed in BRCA1-deficient cells, where CST deficiency confers PARPi resistance (47,48). BRCA1-deficient cells are characterized by impaired DNA end resection, a rate-limiting step in HR, and increasing evidence shows that PARPi resistance in BRCA1-deficent cells arises from restoration or enhancement of end resection (47,48,81–84). Multiple studies have shown that CST restricts end resection at DSBs by inhibiting nuclease attack and/or promoting POLα fill-in (39,47), and its depletion therefore promotes more extensive end resection at the earliest stage during DSB repair, thereby shifting the balance toward hyper-resection. As BRCA1-deficient cells still have functional BRCA2, such increased end resection partially restores HR and reduces reliance on PARP-mediated repair, leading to PARPi resistance. In contrast, BRCA2-deficient cells have intact end resection but are defective at the RAD51 loading step. Therefore, PARPi resistance in BRCA2-deficient cells is expected to arise from the restoration of RAD51 loading. Since CST facilitates RAD51 loading to stalled forks (43), CST overexpression rather than depletion is required for compensating for the RAD51 loading defect and fork deprotection. These opposing effects underscore that CST’s function in modulating PARPi response is highly dependent on whether the primary defect lies in end resection (BRCA1 loss) or RAD51 loading (BRCA2 loss).

Our findings have several important implications. First, STN1 expression level may predict which tumors have acquired alternative mechanisms to protect replication forks. Given the important role of stalled fork protection in PARPi response, expression levels of STN1 could help guide clinical decisions regarding PARPi monotherapy versus combination approaches. Second, targeting the CST-RAD51 interaction or the downstream pathways through which CST stabilizes forks could provide a therapeutic strategy to overcome or prevent resistance. Third, it is possible that additional BRCA2-independent pathways for RAD51 loading may exist and could be exploited by tumor cells under therapeutic pressure. Dissecting these pathways may reveal novel vulnerabilities not only in PARPi-resistant tumors but also in cancers that rely on replication stress tolerance mechanisms more broadly.

Despite the insights gained, several questions remain. It will be important to determine whether STN1 overexpression is driven by genomic amplification, transcriptional upregulation, stabilization of the STN1 protein, or selective outgrowth of STN1-high cells under the PARPi treatment pressure. Further investigation into the regulatory mechanisms controlling STN1 expression will be important. Additionally, because BRCA2 functions in other genome maintenance pathways beyond fork protection such as promoting efficient DNA replication, replication restart, or recombination fidelity, future studies are needed to assess whether STN1 overexpression affects these BRCA2-related processes.

In summary, our findings reveal a novel mechanism of PARPi resistance in BRCA2-deficient cancer cells driven by STN1 upregulation. By restoring RAD51 at stalled replication forks and preventing pathological fork degradation and ssDNA gap accumulation, increasing STN1 abundance likely compensates for BRCA2 loss. These results broaden our understanding of replication stress tolerance pathways in HR-deficient cancers and highlight STN1 as a potential biomarker for PARPi resistance.

## Author contributions

ZAL, NL, SG performed experiments and data analysis. ZAL assembled figures. WC conceived and supervised the study. Q-EW and WC supervised research staff. ZAL and WC wrote the manuscript. WC performed the final editing of the manuscript.

## Acknowledgments

We sincerely thank Dr. George-Lucian Moldovan at Pennsylvania State University Cancer Institute for sharing the HeLa BRCA2 KO cell line, Dr. Kenneth Nephew at Indiana University School of Medicine for providing PEO4 cells, and S. Knowles and R. Jaiswal for technical assistance.

## Data availability

The RNA-seq data generated in this study have been deposited in the Gene Expression Omnibus (GEO) at NCBI with the accession GSE315090. Upon acceptance of the manuscript, the dataset will become publicly available.

