## Supplemental Figures and methods for "STN1 upregulation promotes PARPi resistance in BRCA2-deficient cancer cells via replication fork protection and suppression of ssDNA gap formation"

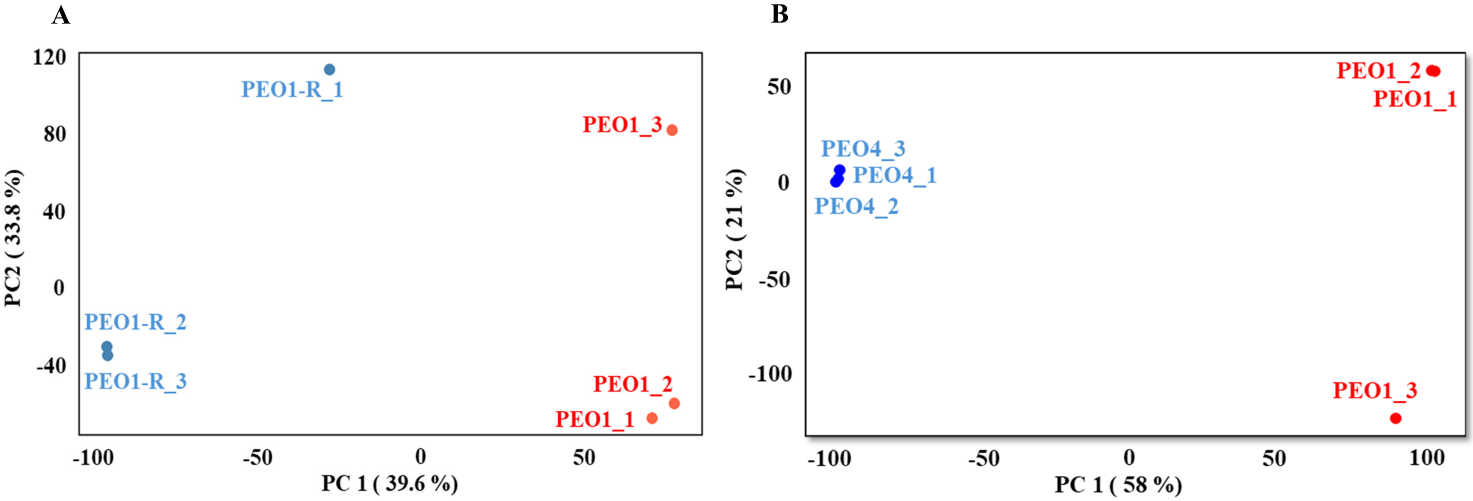


**Supplementary Fig. S1. Transcriptomic profiling reveals distinct gene expression patterns among PEO1, PEO1-R, and PEO4 cells.**

**(A)** Principal component analysis (PCA) of RNA-seq data from PEO1 and PEO1-R cells. Each point represents an individual biological replicate (n = 3 per group), with PEO1 samples shown in red and PEO1-R samples in blue. Principal component 1 (PC1) accounts for 39.6% of the total variance, and principal component 2 (PC2) accounts for 33.8%, demonstrating clear separation between the two cell lines.

**(B)** PCA of RNA-seq data from PEO1 and PEO4 cells. Each point represents an individual biological replicate (n = 3 per group), with PEO1 samples shown in red and PEO4 samples in blue. PC1 accounts for 58.0% of the total variance, and PC2 accounts for 22.0%, demonstrating distinct clustering of the two cell lines.

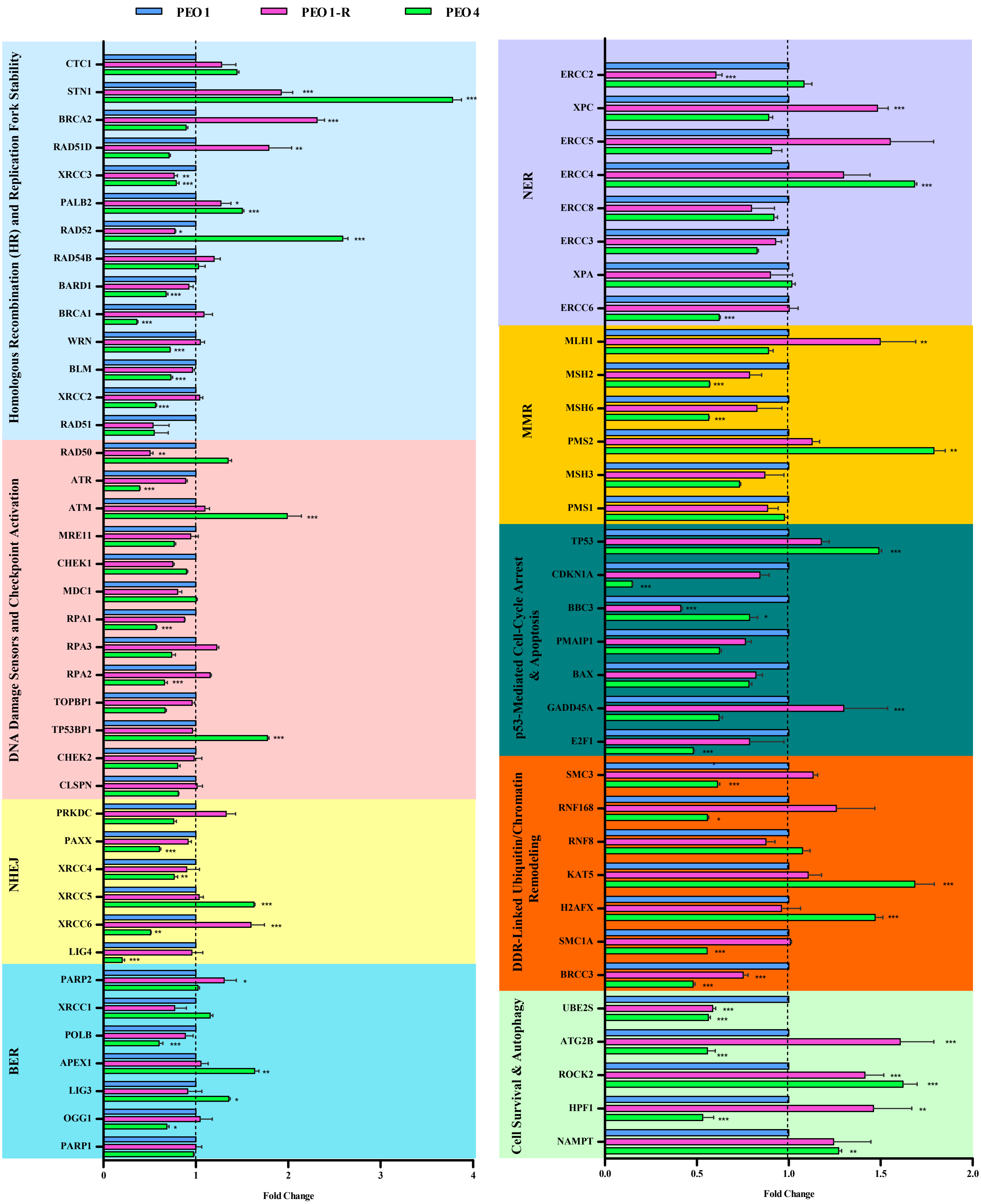


**Supplementary Fig. S2. Transcriptomic profiling reveals differential expression of genes involved in DNA damage response, DNA repair, and cell survival pathways in PEO1, PEO1-R, and PEO4 cells.** Bar graphs showing RNA-seq-derived transcript levels of representative genes involved in DNA damage sensing, DNA repair, and cell survival pathways that might affect the PARPi resistant development and cell survival. PEO1 cells served as control. One-way ANOVA with Tukey's multiple comparisons was used to determine the significance. (* $P < 0.05$, ** $P < 0.01$, *** $P < 0.001$).

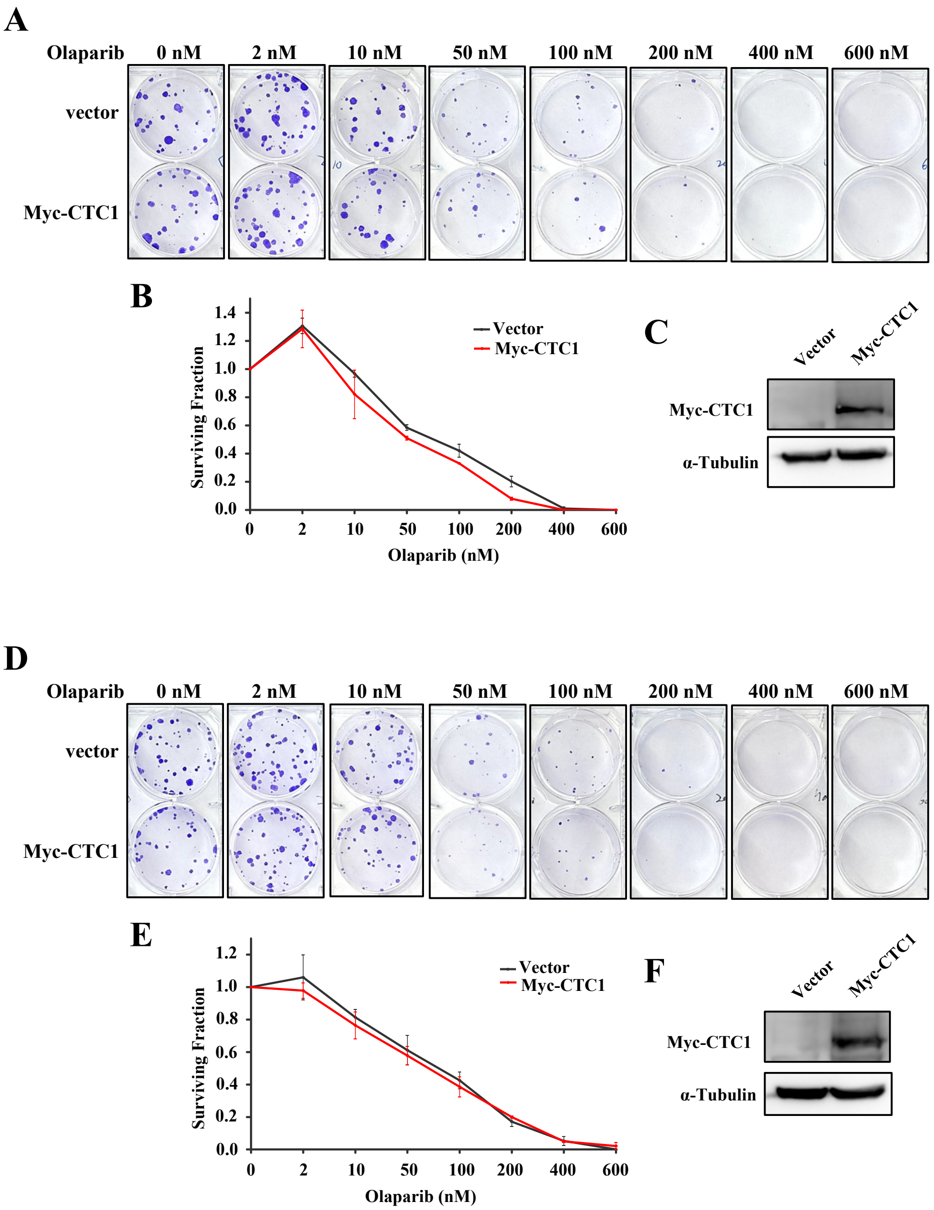


**Supplementary Fig. S3. Effect of ectopical expression of CTC1 on the development of Olaparib resistance.**

**(A)** Representative images of clonogenic assay of PEO1 expressing vector or Myc-CTC1 via retroviral transduction after treatment with indicated concentrations of Olaparib or DMSO. Quantification of survival is shown in **(B).** In all survival curves, data are mean ± SEM of at least two independent experiments. One-way ANOVA with Tukey's multiple comparisons was used to determine the significance. ($^*P < 0.05$)**.**

**(C)** Western blot shows the expression of Myc-CTC1 in PEO1 cells.

**(D)** Representative images of clonogenic assay of PEO1 cells transiently transfected with vector and Myc-CTC1 and treated with indicated concentrations of Olaparib. Quantification of survival after CTC1 overexpression is shown in **(E)**.

**(F)** Western blot shows Myc-CTC1 expression in PEO1 cells.

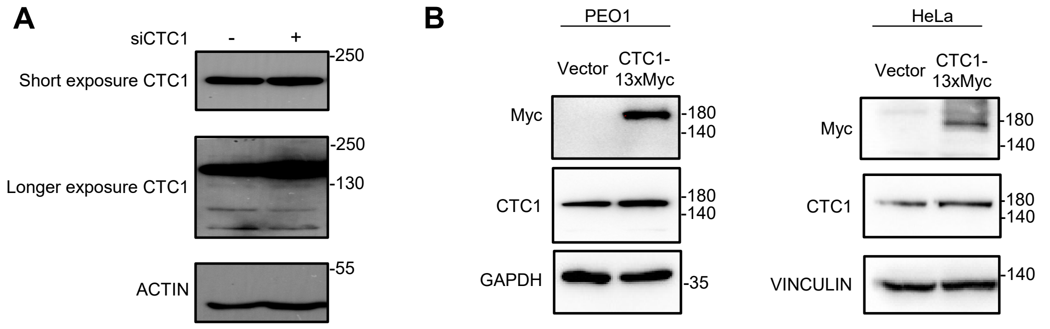


**Supplementary Fig. S4**: **The CTC1 antibody fails to detect CTC1 by western blot.**
**(A)** HeLa cells were transfected with control or CTC1 siRNA, and whole-cell lysates were analyzed by western blot using the CTC1 antibody. Although the antibody detects a specific band, its intensity is not reduced following CTC1 knockdown. Furthermore, the detected band migrates at a substantially higher molecular weight than the predicted size of CTC1 (~130 kDa).
**(B)** The CTC1 antibody fails to detect Myc-tagged CTC1. PEO1 (left) and HeLa (right) cells stably expressing either empty vector or CTC1-13×Myc were analyzed by western blot. An anti-Myc antibody detects CTC1-13×Myc at approximately 170-180 kDa, consistent with previous observation (Lyu et al. EMBO J, 2021). In contrast, the CTC1 antibody does not detect the CTC1-13×Myc protein at this molecular weight.

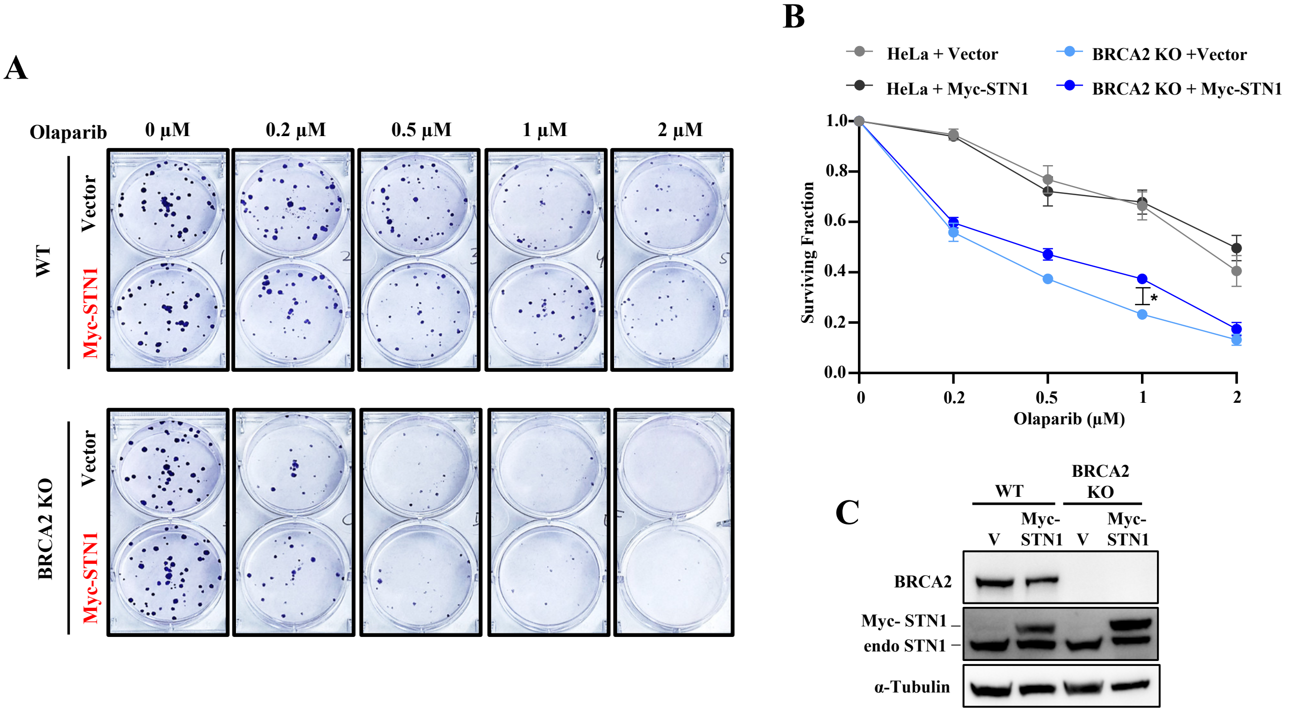


**Supplementary Fig. S5. Elevated STN1 expression increases Olaparib resistance in BRCA2-deficient HeLa cells.**

**(A)** Clonogenic survival of WT and BRCA2 KO HeLa cells with and without Myc-STN1 overexpression. Colonies were stained at day 11 after seeding. Quantification of survival (mean ± SEM from two independent experiments) is shown in **(B)**. *P* values: One-way ANOVA with Tukey's multiple comparisons. $^{*}P < 0.05$.

**(C)** Western blot analysis of BRCA2 expression and elevated STN1 expression in WT and BRCA2 KO HeLa cells transfected with Myc-STN1 or vector.

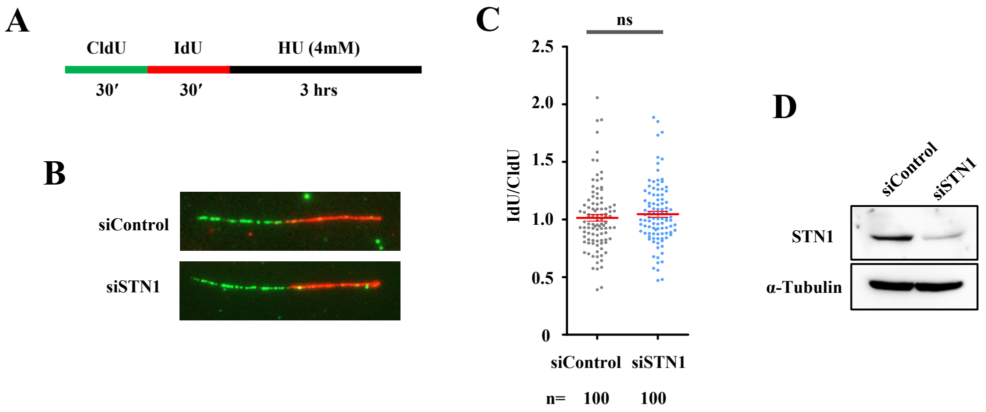


**Supplementary Fig. S6. PEO1-R is proficient in fork protection, and STN1 depletion in PEO1-R does not cause fork degradation.** PEO1-R cells with or without STN1 siRNA were subject to DNA fiber analysis following HU treatment to analyze the degradation of nascent strand DNA.

**(A)** Scheme of CldU-IdU labeling.

**(B)** Representative DNA fiber images.

**(C)** Scatter plots showing the IdU/CldU ratio in PEO1-R cells transfected with control siRNA and siSTN1 followed by HU treatment. The graphs are representative of two independent experiments. n represents the number of DNA fibers analyzed in each sample in each experiment. Statistical significance was determined by Mann-Whitney test.

**(D)** Western blot confirming depletion of STN1 in PEO1-R cells following siRNA transfection.
